# The relationship between multisensory associative learning and multisensory integration

**DOI:** 10.1101/2020.08.28.272633

**Authors:** Sébastien Á. Lauzon, Arin. E. Abraham, Kristina Curcin, Ryan A. Stevenson

## Abstract

Our perception of the world around us is inherently multisensory, and integrating sensory information from multiple modalities leads to more precise and efficient perception and behaviour. Determining which sensory information from different modalities should be perceptually bound is a key component of multisensory integration. To accomplish this feat, our sensory systems rely on both low-level stimulus features, as well as multisensory associations learned throughout development based on the statistics of our environment. The present study explored the relationship between multisensory associative learning and multisensory integration using encephalography (EEG) and behavioural measures. Sixty-one participants completed a three-phase study. First, participants were exposed to novel pairings audiovisual shape-tone pairings with frequent and infrequent stimulus pairings and complete a target detection task. EEG recordings of the mismatch negativity (MMN) and P3 were calculated as neural indices of multisensory associative learning. Next, the same learned stimulus pairs presented in audiovisual as well as unisensory auditory and visual modalities while both early (<120 ms) and late neural indices of multisensory integration were recorded. Finally, participants completed an analogous behavioural speeded-response task, with behavioural indices of multisensory gain calculated using the race model. Significant relationships were found in fronto-central and occipital areas between neural measures of associative learning and both early and late indices of multisensory integration in frontal and centro-parietal areas, respectively. Participants who showed stronger indices of associative learning also exhibited stronger indices of multisensory integration of the stimuli they learned to associate. Furthermore, a significant relationship was found between neural index of early multisensory integration and behavioural indices of multisensory gain. These results provide insight into the neural underpinnings of how higher-order processes such as associative learning guide multisensory integration.

## Introduction

How we process sensory information shapes the manner with which we perceive the world around us, and how we interact with the world. While information from each of sensory modality is transduced independently, it is often integrated into a single, unified perception. Multisensory integration confers a number of behavioural benefits including faster and more accurate perception and behavioural responses (Stein and Meredith, 1993). To reap these benefits, our perceptual systems must perceptualy bind the information that comes from the same external source and segregate the sensory information that come from different sources.

To solve this binding problem, our perceptual systems use two categories of information, lower-level sensory features such as spatial and temporal alignment, and higher-level information such as learned associations and semantic congruence. Semantic congruence (e.g., contextual cues) are often recruited, when pertinent, in multimodal situations, from high-level (Calvert, Campbell et al., 2000) to low-level stimuli (Laurienti, Wallace et al., 2003). More crucially for this experiment, learned multisensory associations play an integral role in whether sensory inputs are bound (Brunel, Carvalho et al., 2015)((Hubel and Wiesel, 1998; Hummel and Gerloff, 2005; Laine, Kwon et al., 2007; Mitchel and Weiss, 2011; Wallace, 2004). As adults, when multisensory stimuli are encountered, there is a tendency to use a combination of stimulus properties, such as temporal synchrony, and previously learned associations (Ten Oever, Sack et al., 2013). These prior experiences are crucial for ensuring accuracy in the interpretation of incoming multisensory information, as the formation of these experiences is complex and multifaceted. These experiences can incorporate semantic, affective, and relational cues into their stored representation, which can make the integration process much more efficient (Lewkowicz, 2014), as top-down effects has been observed as early as 60 ms when exposed to multisensory stimuli (De Meo, Murray et al., 2015). This process changes with age, where infants rely more heavily on the inherent stimulus characteristics than on statistical probabilities of co-occurrence and learned associations when deciding whether to integrate or segregate sensory information (Murray, Lewkowicz et al., 2016). Throughout development, there is a shift from primarily using stimulus features to using learned associations and prior experiences with the world when deciding whether to integrate, a process termed multisensory perceptual narrowing (Lewkowicz, 2014).

This learning of associations between inputs across different sensory modalities can be explained by statistical learning (Sarmiento, Matusz et al., 2016), where statistical regularities are extracted across time in order to learn about the structure of the sensory inputs (Saffran, Aslin et al., 1996). The robustness of this effect can be experimentally demonstrated by presenting participants with novel spatially and temporally congruent audiovisual stimuli that are arbitrarily paired. Over time, participants demonstrated neural and behavioural benefits, in concordance with learning effects (Altieri, Stevenson et al., 2015). It should be noted that such learned associations are distinct from semantic congruency, which is also a top-down process that modulates multisensory integration (Doehrmann and Naumer, 2008).

Learned associations and the stimulus-driven influences on integration do not occur in isolation, but are interactive. Studies have shown that experience with learned associations and their statistics can reduce the strength of temporal factors (Ten Oever, Sack et al., 2013; Habets, Bruns et al., 2017). These findings speak to the constant balance and re-weighting of the pre-attentive, stimulus-driven processes such as temporal and spatial congruence and higher-order processes such learned associations.

Though there is clear theoretical work supporting the link between learned associations across modalities and multisensory integration, to date there have been few studies empirically exploring the relationship between learning novel multisensory associations and how well we integrate information from these associations. Given that associative learning plays a key role in effective integration of sensory information (Murray, Lewkowicz et al., 2016), and that this integration process has been continually associated with behavioural benefits, we posit that multisensory associative learning should then be positively related to multisensory gain. Here, we address this research question by exposing adults to novel audiovisual stimulus pairings (shapes and tones) in a learning phase, and subsequently presenting them with unisensory and multisensory versions of these learned stimulus pairs in a classic multisensory paradigm. We concurrently recorded event-related potentials (ERP) as neural indices of associative learning and multisensory integration, with the prediction that indices of associative learning would be positively related to measures of multisensory integration.

To assess multisensory associative learning, we used a three-stimulus oddball detection paradigm (Courchesne, Hillyard et al., 1975) that included frequent stimuli, infrequent stimuli difficult to discriminate from the frequent stimuli, and a distracter stimulus, which is easily discriminable and highly salient. This version of the oddball task controls for novelty effects to isolate learning (Polich and Comerchero, 2003). Critically, for the three-stimulus oddball detection task, the audiovisual *pairings* comprised the standard, target, and deviant stimuli, as opposed to the unisensory component themselves (Rohlf, Habets et al., 2017). Differences in amplitudes between conditions of interest will be extracted from a *a priori* latency windows. To quantify associative learning, two measures at different latencies will be extracted. The first is the mismatch negativity (MMN; (Näätänen, 1995; Näätänen, Paavilainen et al., 2007), which is a measure of pre-attentive deviance detection that typically occurs in the auditory cortex (Huotilainen, Winkler et al., 1998). The second component is the later going P3b, which has been shown to be representative of potentially inhibitory and encoding processes, and is thought to have parietal and frontal neural generators (Polich, 2007).

Assessing multisensory integration will be achieved using passive exposure to the learned combinations of audiovisual stimuli as well as their unisensory components, while attention is sustained using an irrelevant detection task (Cappe, Thut et al., 2010). Electrophysiological indices of multisensory integration can take place at multiple latencies after stimulus presentation. The first of these indices represents *early* sensory interactions. Such interactions are typically defined as occurring <100 ms post-stimulus onset (De Meo, Murray et al., 2015; Giard and Peronnet, 1999; Foxe, Morocz et al., 2000; Molholm, Ritter et al., 2002), and are typically centrally or fronto-centrally located on the scalp (Talsma, Doty et al., 2007). The second index (approximately 200 ms post-stimulus presentation) represents a later-going index of integration that has been previously established (Besle, Bertrand et al., 2009; Besle, Fort et al., 2005; Giard and Peronnet, 1999). Its topographical scalp locations tend to be over the central, parietal, and occipital areas (Möttönen, Schürmann et al., 2004), and it is thought to be representative of the latest possible latency before confounds such as common activity, which is typically indicative of response selection or motor responses, appear (Besle, Fort et al., 2004; Hillyard, Teder-Salejarvi et al., 1998). Both of these time-windows are thought to represent sensory-perceptual activity that occurs as a result of feedforward bottom-up processes (Foxe, Morocz et al., 2000; Lamme and Roelfsema, 2000), although evidence exists that argues otherwise (Talsma and Woldorff, 2005). Given the passive nature of the stimuli being presented, the audiovisual signal is expected to be subadditive, which represents interactive processes between sensory modalities (Talsma, Doty et al., 2007; Stevenson, Ghose et al., 2014; Hein, Doehrmann et al., 2007; Vroomen and Stekelenburg, 2011).

Finally, a follow-up behavioural measure of multisensory integration will be used (with the same stimuli as is used in the rest of the experiment) as a validation measure for use in quantifying multisensory integration. It will also be compared to the measures of multisensory associative learning.

## Methods

### Participants

Participants were 65 undergraduate students aged 17-55 at the University of Western Ontario. Four participants were excluded as they failed to complete the experiment (4 female, 4 right handed). The final sample included *N* = 61 participants (21 males, 4 left-handed) participants aged 17 to 55 years (*M* = 18.97, *SD* = 5.27). Participants completed three computer tasks. The first part of the study was a multisensory associative learning task, and the second a multisensory integration task, both wherein electroencephalographic (EEG) activity was recorded at the scalp. The last part of the experiment consisted of a behavioural measure of multisensory integration.

### Equipment

Electrophysiological data were collected using a 128-channel Hydrocel GSN EGI (Electrical Geodesics Inc., Eugene, OR, USA) cap and sampled at a rate of 250 Hz. All visual stimuli were presented on an LCD screen for the EEG components, and on a CRT screen for the behavioural component to collect precise response times, both with a 60 Hz refresh rate. All auditory stimuli were presented via a speaker on either side of the participant, 160 cm from their head. Responses were collected using a Serial Response Box (Model 200A; Psychology Software Tools, Inc., 2003). Experiments were conducted using E-Prime 2.0.8.252. (Psychology Software Tools, Inc., 2014) using NetStation Extensions version 2.0. The experiment took place in a sound-attenuated booth (background dB SPL = 30.4 dB).

### Stimuli

Auditory stimuli consisted of pure tones created using Matlab’s Psychophysics Toolbox (Kleiner, Brainard, & Pelli, 2007). The frequencies of the tones were chosen to ensure adequate perception and discriminability. The three tones of distinct frequencies (320.00 Hz, 427.15 Hz, and 570.14 Hz), were 100 ms in duration, were sampled at a rate of 8000 Hz, and played at 82-83 dB SPL. The auditory features will be referred to as A1, A2, and A3.

Visual stimuli were presented through a computer screen on a black background. Visual stimuli were three white two-dimensional shapes (circle, square, and triangle) presented on a black background, and created using Adobe Illustrator CC. The shapes were controlled for luminance by keeping their area constant. The visual angles (width x height) of the circle, square, and triangle were 8.86° x 8.86°, 7.82° x 7.82°, and 11.89° x 10.38°, respectively. These visual features will be referred to as V1, V2, and V3.

### Procedure

#### Phase 1: Multisensory Associative Learning Phase

Throughout this phase, participants were presented with audiovisual tone-shape pairings, each pair with its own frequency of presentation (see Table 1 for a complete layout of presentation frequencies). Participants were tasked with responding with their right index finger, by using the serial response box, as quickly and as accurately as possible to a specific audiovisual pairing, “Target”. Two pairings, A1V1 and A2V2, were presented during 70% of total trials (35% each), and will subsequently be referred to as “Match” trials. A1V2 pairings were presented on 10% of trials, and will be referred to as “Mismatch” trials. A2V1 pairings were also presented on 10% of trials, and were target trials to which participants were instructed to respond. Finally, the A3V3 pairing was presented for 10% of trials, and will be referred to as “Deviant” trials. Deviant trials were included in order to control for attention-switching due to rare sensory features (Rohlf, Habets et al., 2017). The three visual stimuli (circle, square, triangle) and three auditory stimuli (high, medium, low), were counterbalanced across participants.

**Table 1:**
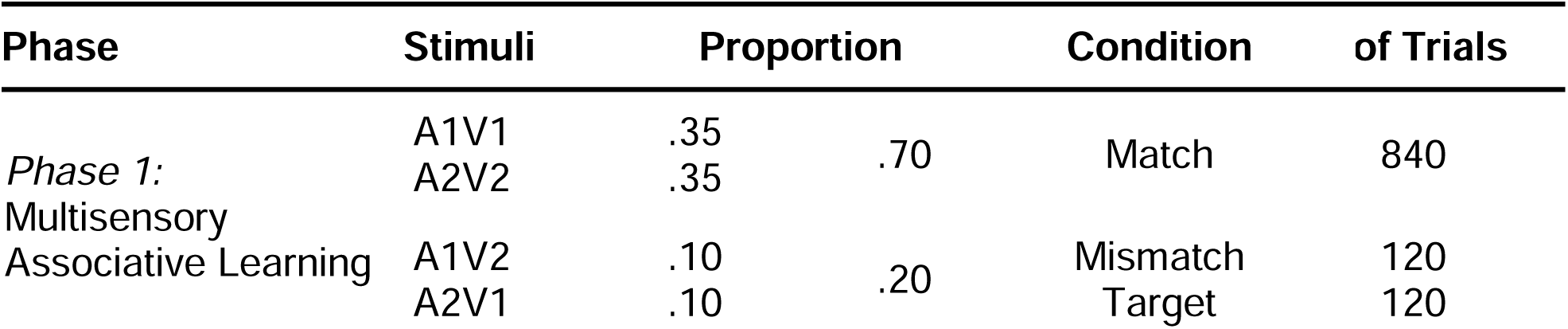
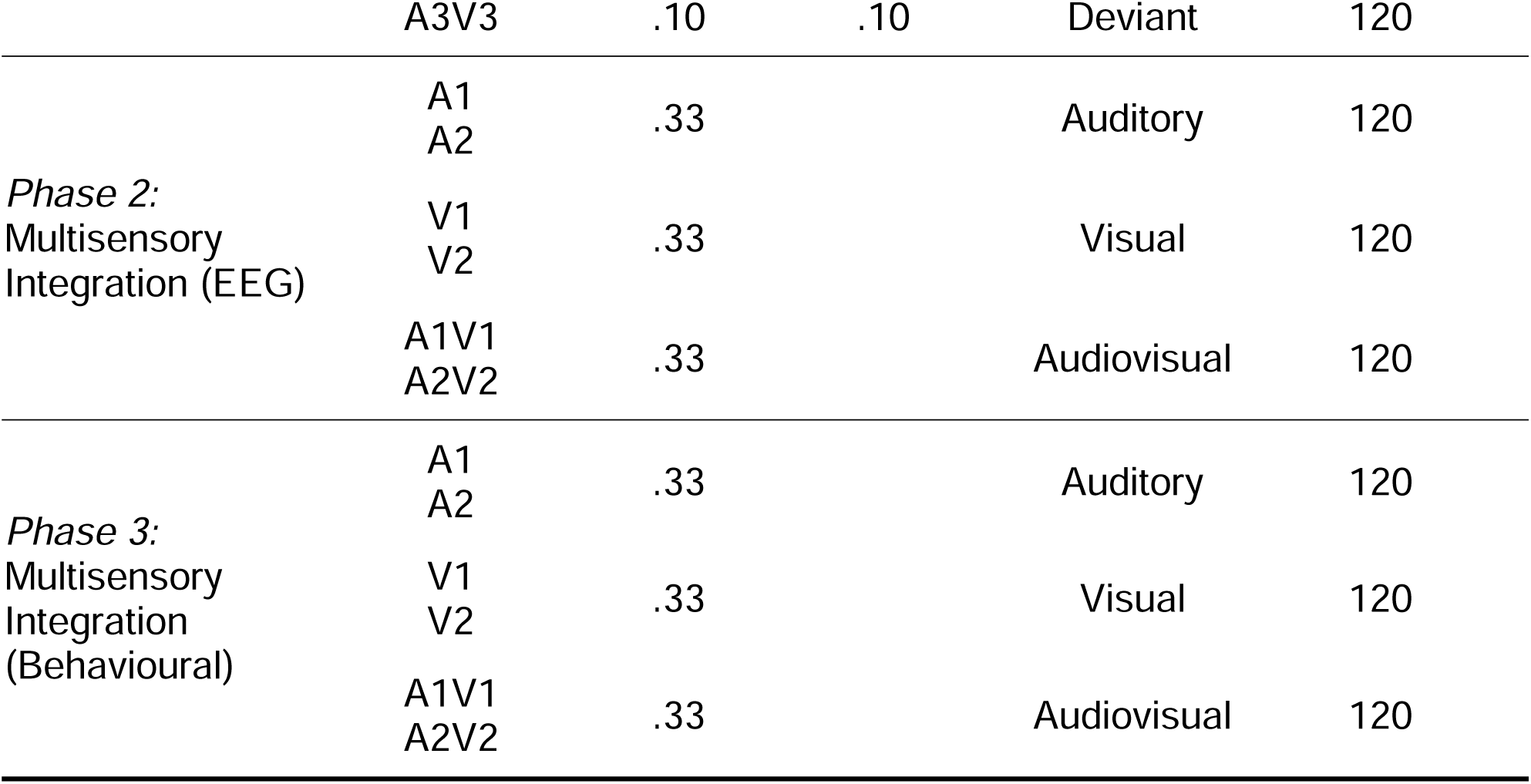
Experimental Design of Phases 1, 2, and 3.

Each trial consisted of a 100 ms audiovisual stimulus presentation followed by an inter-trial interval where a white visual fixation cross was shown for a randomly jittered duration of 900-1400 ms. At the beginning of the experiment, participants were instructed to respond by pressing the leftmost button on a serial response box (‘1’) when they detected the target combination which was presented to them immediately prior to testing (Figure 1A). Responses were recorded during the inter-trial interval where the white fixation cross was presented. This phase of the experiment was comprised of a total of 1200 trials, which were presented in random order, and divided into five blocks of 240 trials with short periods of rest to check the impedances on the EEG net. Thus, a total of 840 match, 120 mismatch, 120 target mismatch, and 120 deviant trials were presented during this phase of the experiment.

**Figure 1.**
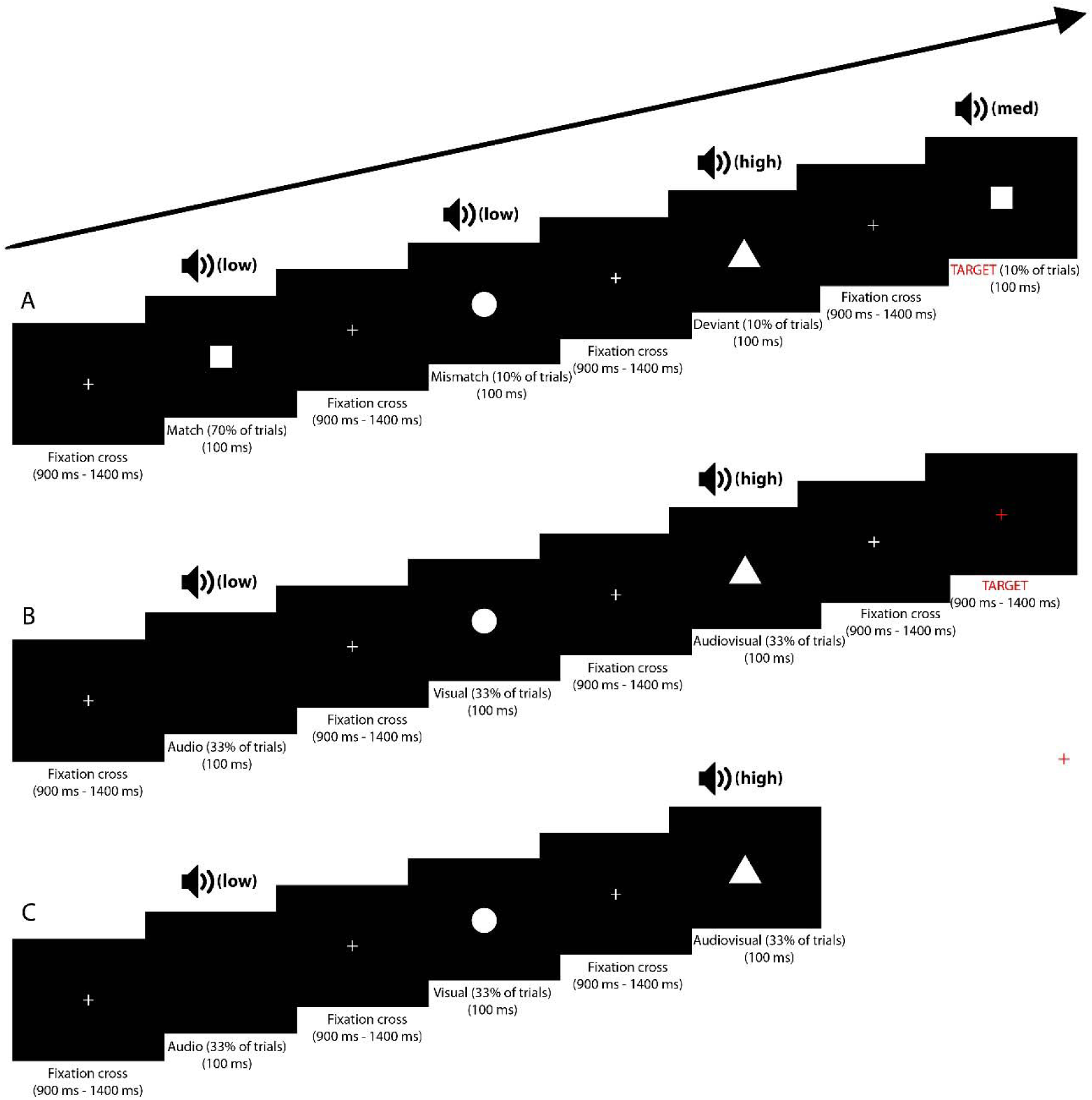
Experimental design. Panels A-C depict trial structures for the ERP associative learning, ERP multisensory integration, and behavioural multisensory integration paradigms, respectively.

#### Phase 2: Multisensory Integration Phase (EEG)

This second phase used the same features of the stimuli from the associative learning phase to test for multisensory integration. Presentations of the visual and auditory unisensory components of the match stimuli were included (A1, A2, V1, V2), as well as matched audiovisual presentations (A1V1 and A2V2). Note that the audiovisual combinations presented in this phase were always the matched, frequently-presented pairings, never the mismatched, target, or deviant stimulus pairs from the previous phase. Trial structures were the same as in the associative learning phase, with the exception that following 10% of trials, the fixation cross turned red 100 ms after the initial fixation presentation. Participants were tasked with responding via key press when this red fixation appeared in order to assure vigilance while not contaminating EEG recordings with a motor artifact during stimulus presentations (Figure 1B). There was a total of 360 trials, which were equally distributed across conditions, 120 audio-only, 120 visual-only, and 120 audiovisual trials. A break was included after 180 trials.

#### Phase 3: Multisensory Integration Phase (Behavioural)

This portion of the experiment tested for a behavioural measure of multisensory integration using the same paradigm as its analogous EEG phase. However, in this portion of the experiment, participants were instructed to respond via response box as quickly as possible when they detect<u>ed</u> either an auditory, visual, or audiovisual stimulus with response times (RTs) recorded. No red fixation cross was presented in this portion of the experiment (Figure 1C).

### Analysis

Data was collected using continuous EEG recording through EGI NetStation, and analyzed using NetStation Waveform Tools and Matlab. Data were initially band-pass filtered at 0.1-100 Hz. Additionally, a 60 Hz notch filter was applied to filter out powerline interference. Only correct trials (correctly identifying the target, and correctly withholding a response for all other trials) were included in the analyses. Epochs of 1200 ms were extracted from the data, with the first 200 ms used for baseline correction, and the last 1000 ms post-stimulus presentation. Epochs in which motion artifacts such as eye blinks (>50 µV, window size = 640 ms; moving average = 80 ms) or eye movements (>50 µV, window size = 640 ms; moving average = 80 ms) were excluded. Bad channels (>150 µV, across entire segment; moving average = 80 ms) were removed based on whether 20% of the segments were identified as “bad”. These channels were replaced by spherical spline interpolating the signal from the surrounding electrodes. An epoch was deemed “bad” if it contained more than 20 bad channels, contained an eye blink, or contained an eye movement. Bad epochs were excluded from analyses. An average reference was computed, and data was re-referenced to the average.

#### Phase 1: Multisensory Associative Learning Phase

For the associative learning phase of the experiment, the MMN and P3b time-windows were defined as time-window latencies observed in previous literature, which were 100-250 ms (Näätänen and Winkler, 1999) and 300-600 ms (Polich and Comerchero, 2003) respectively. Within these *a priori time windows*, latencies were identified where there were five consecutive time points showing a significant amplitude difference between the match and mismatch conditions for individual participants’ waveform, tested with a paired-sample t-test (α = .05 for each time point). Within these significant time-windows, *a priori* defined electrode clusters that outline anatomical regions of the brain (Tripathi, Mukhopadhyay et al., 2018) were extracted. Clusters with multiple electrodes showing significant amplitude differences for the MMN and P3b were used in the analysis. Significant electrodes contiguous with a predefined cluster with multiple significant electrodes were incuded in this cluster, given that they were not already assigned to a predefined cluster of activity with multiple significant electrodes.

The mean amplitude of these significant windows was used to quantify multisensory associative learning, as mean relative to peak amplitude is less sensitive to noisy data and is effective whenever the latency windows are well established (Luck and Gaspelin, 2017). Both MMN and P3b values were calculated for each individual by subtracting the match from the mismatch mean values within their respective time windows.

Participants’ data were considered outliers if their mean difference scores between the conditions of interest were more than three times the value of the interquartile range for an electrode cluster at either the early or late time window. Data from participants who were outliers were imputed using a Markov Chain Monte Carlo multiple imputation with a maximum of 100 iterations. Imputations were conducted 10 times, with the mean value of these 10 imputations used.

#### Phase 2: Multisensory Integration Phase (EEG)

For the multisensory integration phase, the amplitudes from the unisensory and multisensory signals were compared to quantify multisensory interactions. As electrical fields detected by EEG sum linearly, interactions between auditory and visual processing are identified by summing the two unisensory signals and comparing this sum to the audiovisual signal, known as the additive criterion (Besle, Fort et al., 2004; Stevenson, Ghose et al., 2014). Interactions are thus defined by significant differences:

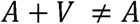

Two windows were extracted based on previous literature, an early (∼40-110 ms) and a late (140-220 ms) latency range of multisensory integration (Giard and Peronnet, 1999; Molholm, Ritter et al., 2002). Criteria for identifying electrodes with significant amplitude differences and for cluster extraction were defined using the same specifications as the previous phase. Values for mean amplitudes were then extracted for both audiovisual presentations and the summed unisensory presentations. The level of multisensory integration was calculated for each individual by subtracting the summed unisensory from the audiovisual values within early and late time windows within each cluster.

Participants’ data were considered outliers if their mean difference scores between the conditions of interest were more than three times the value of the interquartile range for an electrode cluster at either the early or late time window. Data from participants who were outliers were imputed using a Markov Chain Monte Carlo multiple imputation with a maximum of 100 iterations. Imputations were conducted 10 times, with the mean value of these 10 imputations used. If a participant was identified as an outlier in both Phase 1 and Phase 2, the participant’s data was removed from analysis in both phases.

#### Phase 3: Multisensory Integration Phase (Behavioural)

The Race Model (Miller, 1982; Raab, 1962) is commonly used to test for behavioural multisensory integration, and postulates that integration could be present if the mean response times from the multisensory stimuli are smaller than that of either of their unisensory components, assuming that the processes do not interact with one another. In this case, the response times from the behavioural multisensory integration phase were compared using the same principle as their EEG counterpart. Cumulative distribution functions (CDFs) of the response times are calculated for each of the unisensory components, and then summed. These represent the predicted response times, assuming independent processing, also known as Miller’s bound (Miller, 1982). The CDF of RTs during audiovisual trials was then computed and compared to Miller’s bound. Violations of Miller’s bound occur when the audiovisual CDF is above and to the left of Miller’s bound, i.e., when RTs in response to audiovisual presentations occur faster than predicted by responses to the unisensory presentations, and are indicative of multisensory integration/facilitation. Otto’s redundant signals effect (RSE) toolbox was used to compute Miller’s bound, as well as the violation values (Otto, 2019). A binomial test was used to assess whether a significant number of individual participants showed multisensory enhancement.

### Relating Learning to Integrating

Bivariate Pearson correlations were performed between the mean MMN and P3b values and the mean of the difference in both early and late MSI windows to determine whether a relationship existed between participants’ multisensory associative learning performance and their multisensory integration abilities across each cluster. Corrections for multiple comparisons were performed by controlling the false discovery rate by using the Benjamini-Hochberg procedure (false discovery rate (Q) = .05) (Benjamini and Hochberg, 1995).

### Relating Behavioural to EEG Multisensory Integration Measures

Bivariate Pearson correlations were also performed between the EEG and behavioural measures of multisensory integration. This analysis was included as a validation measure for the EEG measure of multisensory integration.

## Results

### Phase 1: Multisensory Associative Learning

An average of 1178.87 trials (98.24% of total trials) per participant were included in the analysis. Excluded trials were both incorrectly identified targets and target misses. For this phase of the experiment, a total of 7 participants’ data was identified as outliers, and scores were imputed for 5 of them. The following analyses for this phase of the experiment therefore include 59 participants.

A cluster exhibiting a significant difference between the Mismatch and Match conditions was found in the left parieto-occipital area (LPO; electrodes 60, 52, 51, 67, 59, 58, 71, 66, 65, 64, 70, 69, 74, and 68) in the MMN latency range, between 216-252 ms (Figure 2A). Significant differences between Mismatch and Match conditions were only found in the left hemisphere, therefore, the right hemisphere was not considered for this measure. The mean amplitude difference between the Match and Mismatch conditions was *M* = .477 µV, *SEM* = .145 µV (Figure 3), which was significant (*t*(58) = 3.296, *p* = .002, *d* = .429). The mean difference between the Deviant and Match conditions *M* = .350 µV, *SEM* = .174 µV, was significant (*t*(58) = 2.004, *p* = .0498, *d* = .261).

**Figure 2:**
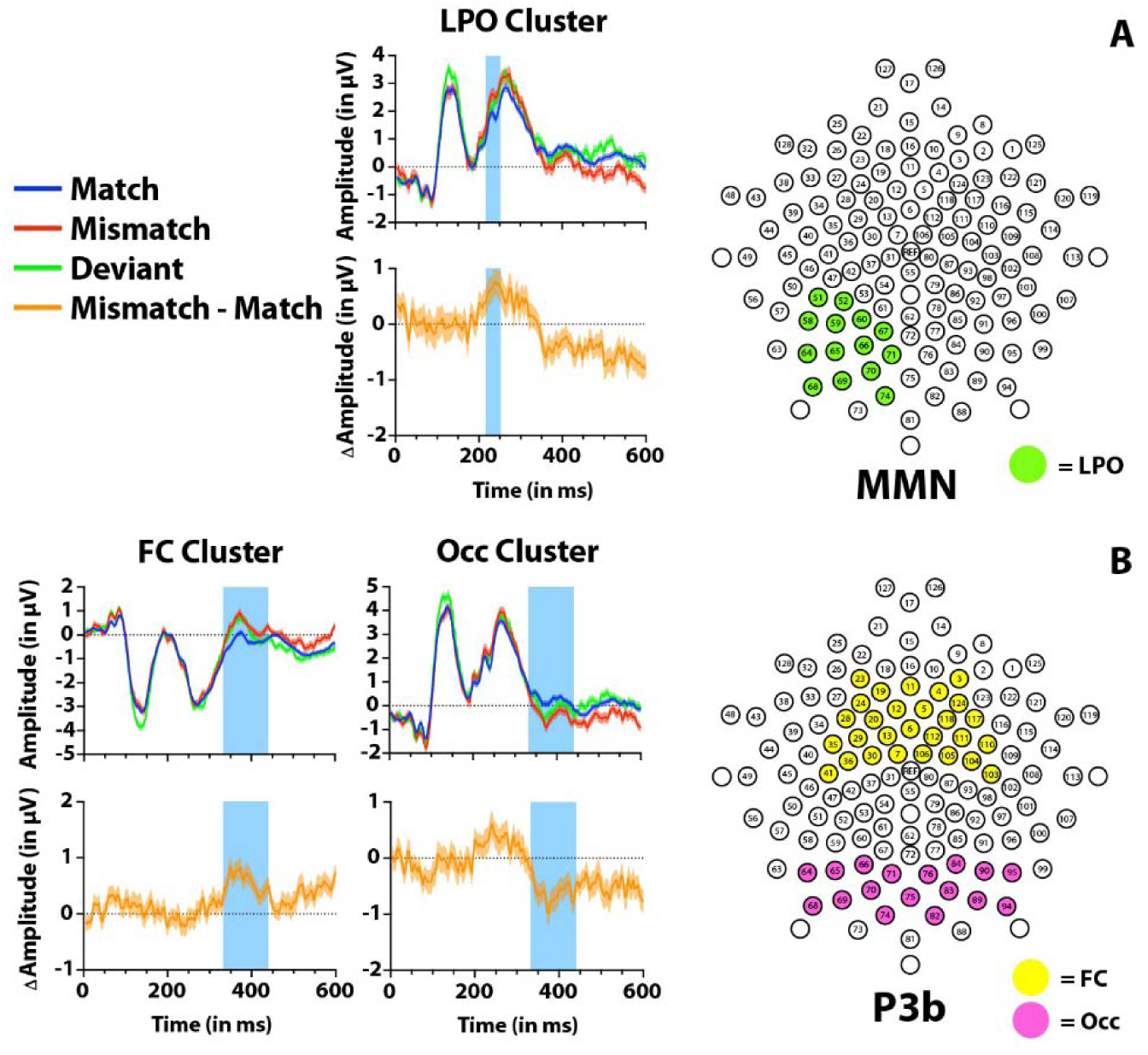
Scalp topography and timecourses for Phase 1, the EEG portion of the associative learning phase. The envelope around the individual time courses represents the standard error of the mean (SEM). The orange timecourse represents the activity from the Match condition subtracted from the Mismatch condition. A), The extracted cluster for the MMN, the left parieto-occipital (LPO) cluster, is portrayed on the right, with the timecourses for the individual conditions on the left. B), The extracted clusters for the P3b, the fronto-central (FC) cluster and the occipital (Occ) cluster are portrayed on the right, with the timecourses for the individual conditions on the left.

**Figure 3:**
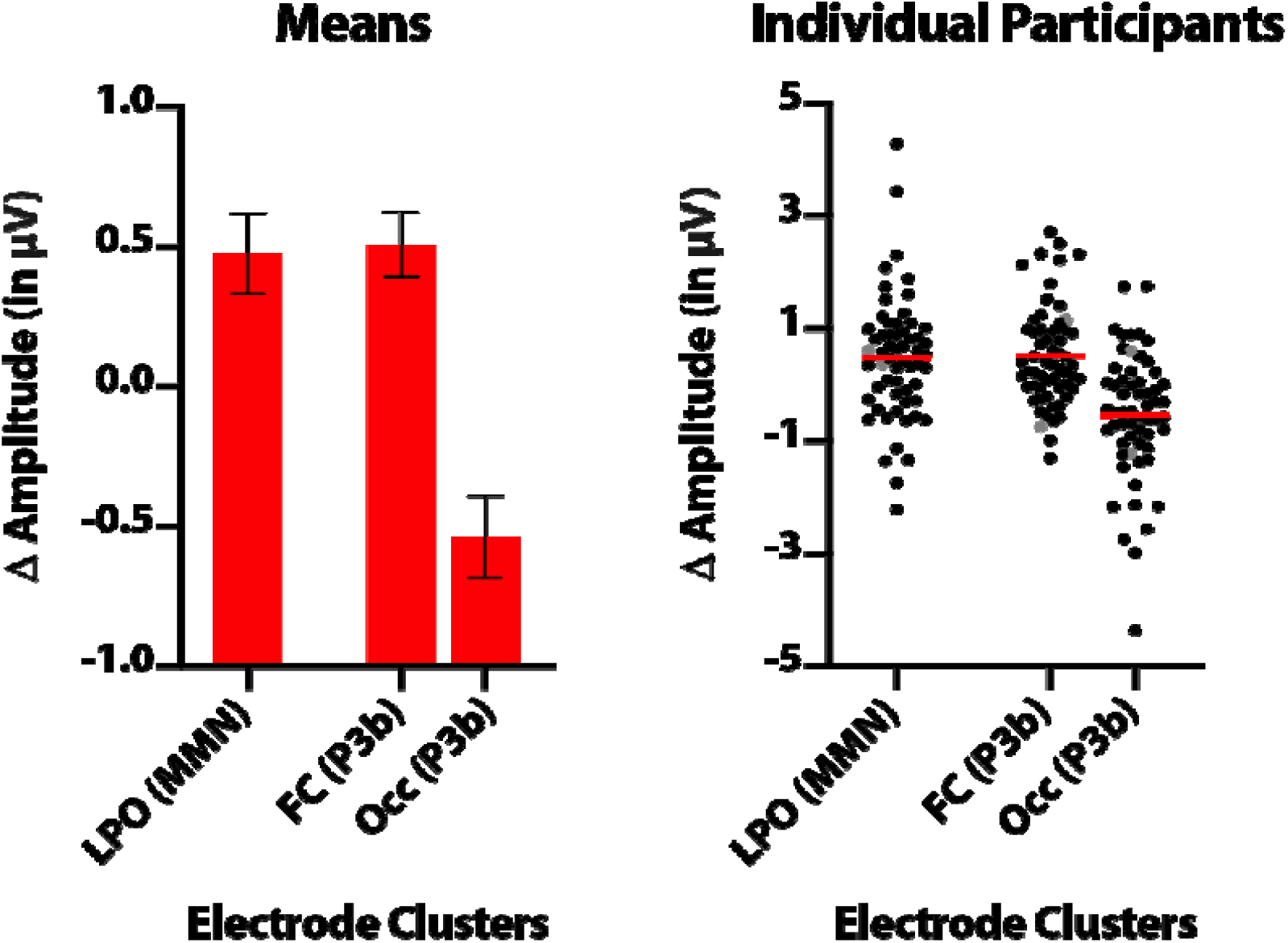
Group means and individual means for the electrode clusters corresponding to each measure of Phase 1, multisensory associative learning. Error bars represent SEM, and the red lines correspond to the mean. The grey individual data points represent participants who are more than 3 SD away from the age mean but were still included in analyses.

For the P3b latency range, between 332-440 ms, the first significant electrode cluster was fronto-central (FC; electrodes 11, 6, 3, 4, 124, 5, 118, 117, 23, 19, 24, 12, 20, 28, 112, 111, 110, 106, 105, 104, 103, 13, 29, 35, 7, 30, 36, and 41; Figure 2B). There were no significant hemispheric differences (*t*(58) = .088, *p* = .930, *d* = .011) and as such, both hemispheres were collapsed into one cluster. The mean difference between the Mismatch and Match conditions was *M* = .509 µV, *SEM* = .116 µV (Figure 3), which was significant (*t*(58) = 4.371, *p* < .001, *d* = .569). The difference in amplitudes between the Deviant and Match trials for this cluster was also significant (*t*(58) = 3.459, *p* = .001, *d* = .450).

In the same latency range, an occipital (Occ; electrodes 84, 76, 90, 95, 83, 89, 82, 94, 75, 71, 66, 65, 64, 70, 69, 74, and 68) (Figure 2B) electrode cluster was also extracted. There were no significant hemispheric differences (*t*(58) = .922, *p* = .360, *d* = .120) and as such, both hemispheres were collapsed into one cluster. There was a mean amplitude difference between the Mismatch and Match conditions of *M* = −.536 µV, *SEM* = .146 µV (Figure 3), which was significant (*t*(58) = 3.658, *p* < .001, *d* = .476). The difference in amplitudes between the Deviant and Match conditions was also significant (*t*(58) = 3.685, *p* < .001, *d* = .478).

### Phase 2: Multisensory Integration (EEG)

An average of 359.88 trials per participant, with a task accuracy rate of 99.97% were included in the analysis for this phase of the experiment. Trials were excluded if they were incorrectly identified as the red fixation cross target, as that data was then contaminated by a motor response. All differences below refer to amplitude differences between the sum of the unisensory conditions (Audio + Visual) and the Audiovisual condition (AV). Four participants’ data were identified as outliers, and following this observation, two of these were imputed. The total number of participants for this phase of the experiment was 59.

A single significant central electrode cluster for the early latency window was identified between 48-100 ms (C; electrodes 106, 105, 104, 80, 87, 93, 7, 30, 36, 55, 31, 37, 42, 79, 86, 92, 98, 97, 78, 85, 77, 91, 76, 84, 54, 53, 47, 62, 72, 61, 60, 52, 51, 67, 59, 71, and 66) (Figure 4A). No significant difference between hemispheres was detected (*t*(58) = −.211, *p* = .833, *d* = .028) and as such, hemispheres were collapsed into a single cluster. A mean difference of *M* = .628 µV, *SEM* = .119 µV (Figure 5) was found to be significant (*t*(58) = 5.289, *p* < .001, *d* = .689), where the Audiovisual condition was subadditive in comparison to the sum of the auditory and visual components.

**Figure 4:**
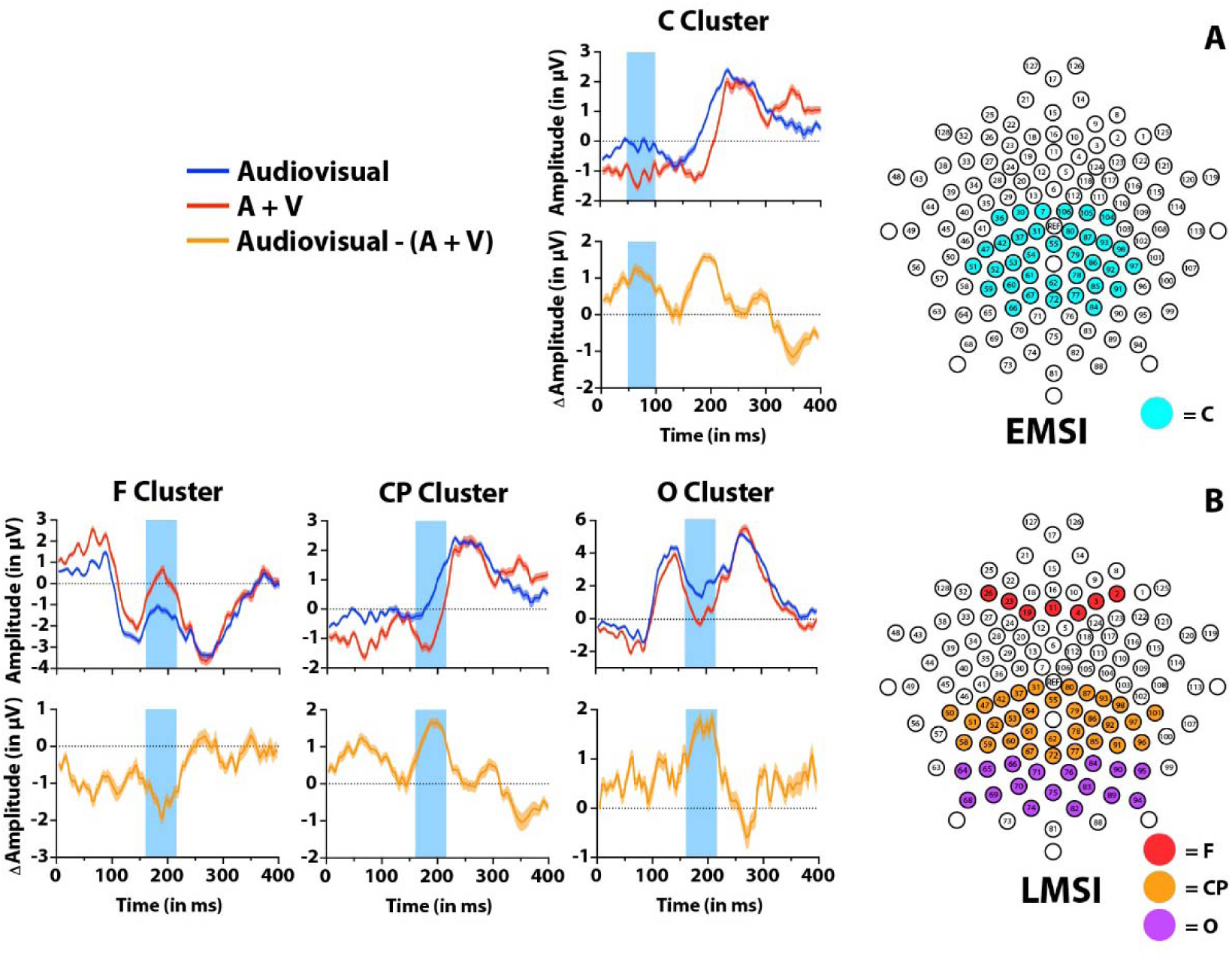
Scalp topography and timecourses for Phase 2, the EEG portion of the multisensory integration phase. The envelope around the individual timecourses represents the standard error of the mean (SEM). The orange timecourse represents the activity from the summed Auditory and Visual conditions (A + V) subtracted from the Audiovisual condition (AV). A), The extracted cluster for the early window of multisensory integration (EMSI), the central (C) cluster, is portrayed on the right, with the timecourses for the individual conditions on the left. B), The extracted clusters for the later window of multisensory integration (LMSI), the frontal (F) cluster, the centro-parietal (CP) cluster, and the occipital (O) cluster are portrayed on the right, with the timecourses for the individual conditions on the left.

**Figure 5:**
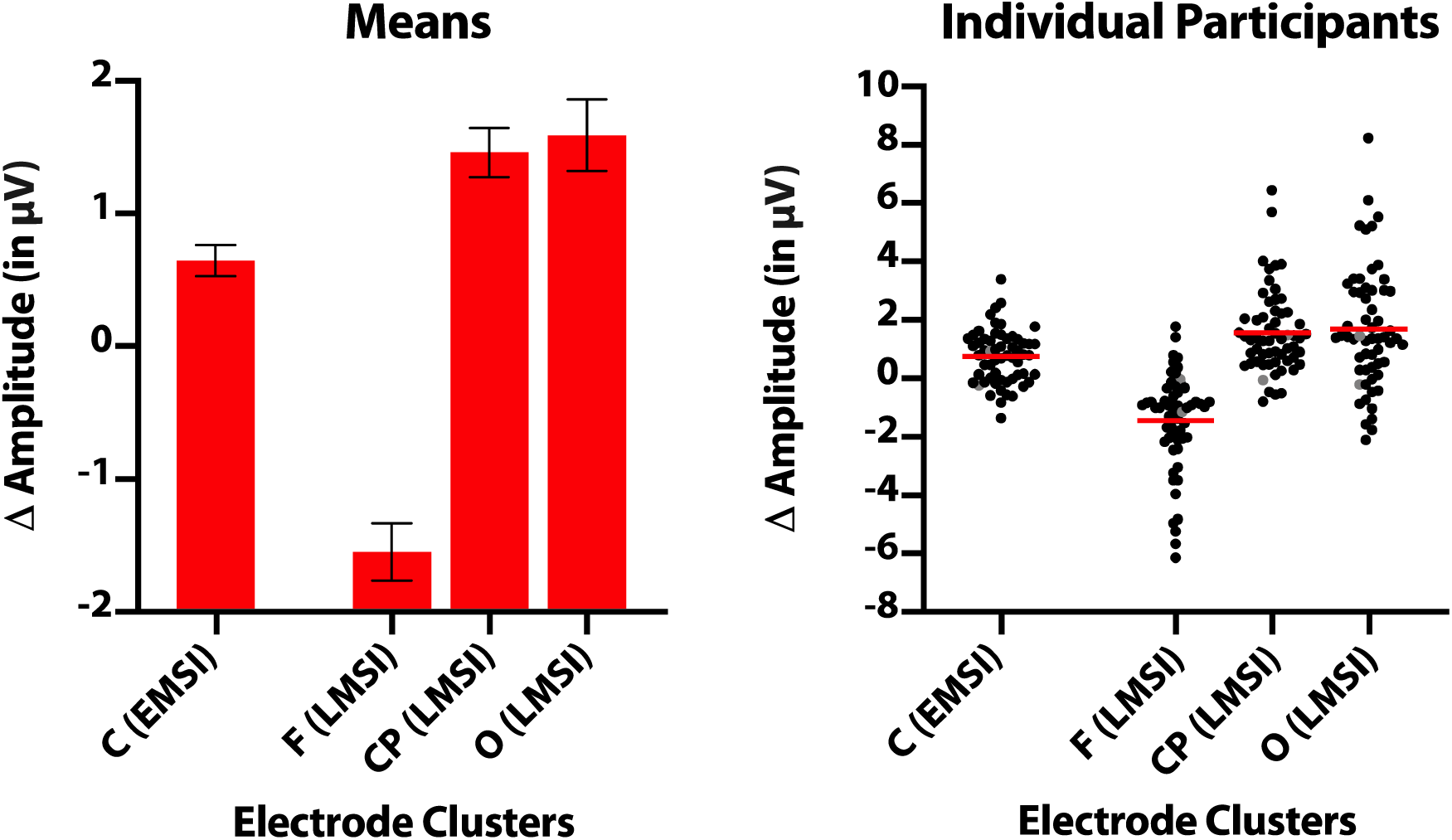
Group means and individual means for the electrode clusters corresponding to each measure of Phase 2, multisensory integration. Error bars represent SEM, and the red lines correspond to the mean. The grey individual data points represent participants who are more than 3 SD away from the age mean but were still included in analyses.

Three significant electrode clusters for a later latency window of 160-216 ms were extracted. A small frontal cluster (F; electrodes 2, 3, 4, 11, 26, 23, and 19) (Figure 4B) showed no significant hemispheric differences (*t*(58) = −.124, *p* = .902, *d* = .016), and as such the data were collapsed across hemispheres. This cluster showed subadditivity, where the amplitudes of the sum of the unisensory components was greater than the audiovisual component, with a mean difference of *M* = −1.571 µV, *SEM* = .216 µV (Figure 5), which was significant (*t*(58) = 7.264, *p* <.001, *d* = .946).

A second, centro-parietal cluster (CP; electrodes 80, 87, 93, 55, 79, 86, 92, 98, 97, 101, 78, 85, 62, 77, 91, 96, 72, 31, 37, 42, 54, 53, 47, 61, 60, 52, 51, 50, 67, 59, and 58) was also extracted (Figure 4B). The cluster collapsed electrodes across hemispheres, as no significant hemispheric differences were detected (*t*(58) = −.784, *p* = .436, *d* = .102). This cluster showed subadditivity, where a difference of *M* = 1.441 µV, *SEM* = .185 µV (Figure 5) was found. This difference was significant (*t*(58) = 7.812, *p* < .001, *d* = 1.017).

A final, occipital cluster (O; electrodes 71, 66, 65, 64, 70, 69, 74, 68, 84, 75, 76, 90, 95, 83, 89, 82, and 94) was extracted (Figure 4B). The electrodes were collapsed across hemispheres, as no significant hemispheric differences were observed (*t*(58) = 1.643, *p* = .106, *d* = .214). This cluster showed subadditive activity, where a difference of *M* = 1.572 µV, *SEM* = .269 µV (Figure 5) was found, which was significant (*t*(58) = 5.848, *p* <.001, *d* = .762).

### Phase 3: Multisensory Integration (Behavioural)

The mean violation of Miller’s bound was *M* = .001, *SEM* = 2.45e-04 (Figure 6A). A binomial analysis revealed that the proportion of participants showing race model (Miller’s bound) violations, in 45 out of 58 participants, was significantly greater than chance (*p* = .000023).

**Figure 6:**
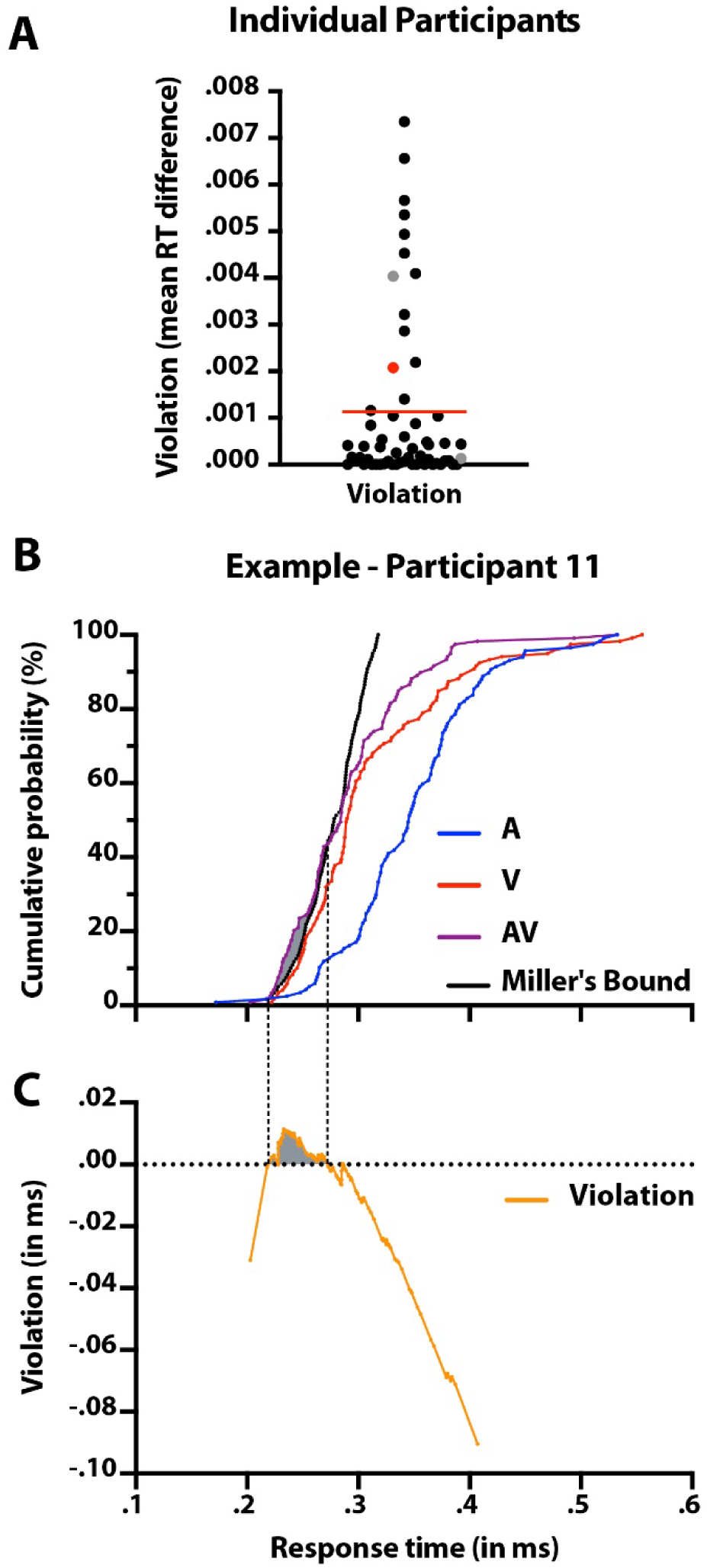
Panel A, Race model violation, representing Miller’s bound violation for individual participants. This value represents the area of the violation or the mean RT difference. The red line represents the group mean and the grey individual data points represent participants who are more than 3 SD away from the age mean but were still included in analyses. The red data point is used as an in Panels B and C. Panels B and C represent an example participant, illustrating the cumulative distribution functions of the multisensory condition as well as both unisensory conditions and Miller’s bound. The violation is represented by the shaded area.

### Relating Learning to Integrating

Early measures of multisensory associative learning (the MMN) in the left parieto-occipital cluster were not significantly correlated with any index of multisensory integration (see Table 2). Conversely, associative learning as measured by the P3b in the fronto-central cluster was significantly correlated to both early multisensory integration in the central cluster (*r*(57) = −.544, *p* = 8.466e-06) (Figure 7A), and later multisensory integration in the centro-parietal scalp area (*r*(57) = −.404, *p* = .001) (Figure 7C). Similarly, the occipital scalp area during later associative learning had a significant correlation between early integration in the central cluster (*r*(57) = .446, *p* = 4.033e-04) (Figure 7B), and later multisensory integration in the centro-parietal scalp area (*r*(57) = .352, *p* = .006) (Figure 7D). All of the significant correlations reported here have been deemed significant using the Benjamini-Hochberg procedure (Benjamini and Hochberg, 1995) with a false discovery rate of Q = .05.

**Table 2:** Correlations – correlation coefficient (*p* value)

| Correlation clusters | EMSI C | LMSI F | LMSI CP | LMSI O | Behavioural MSI |
| --- | --- | --- | --- | --- | --- |
| MMN LPO | -.199 (.130) | .162 (.219) | -.231 (.079) | -.097 (.463) | .083 (.537) |
| P3b FC | **.544 (8.466e-06\*\*)** | -.085 (.521) | **.404 (.001\*)** | .046 (.727) | -.092 (.491) |
| P3b Occ | **-.446 (4.033e-04\*\*)** | .115 (.387) | **-.352 (.006\*)** | -.032 (.809) | .123 (.357) |
| Behavioural MSI | **-.322 (.014\*)** | .086 (.522) | -.070 (.603) | -.063 (.640) | -- |

Note: \* $p < .05$ , \*\* $p < .01$

**Figure 7:**
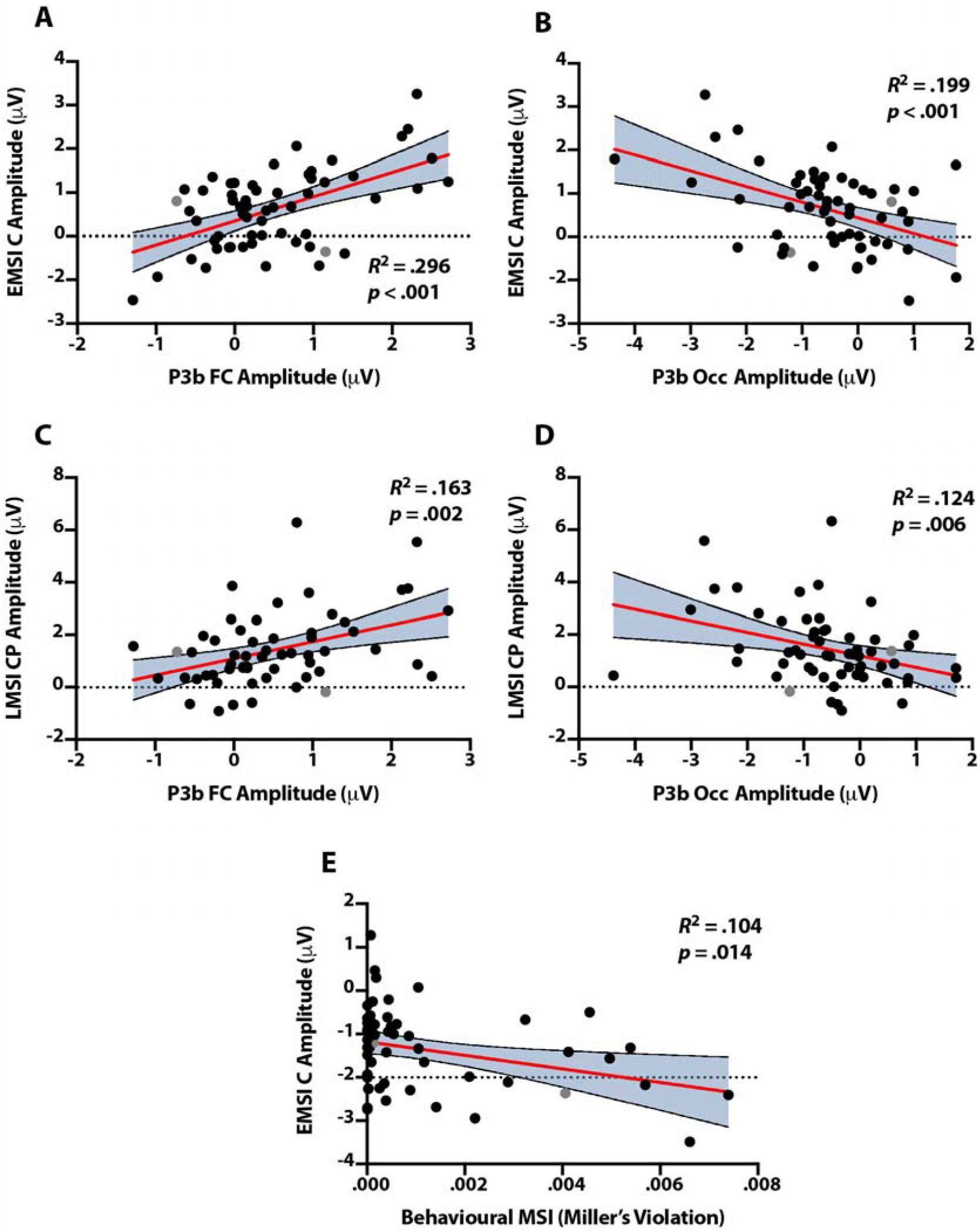
Significant correlations of interest with a 95% confidence interval envelope around the regression line. The grey individual data points represent participants who are more than 3SD away from the age mean, but were still included in analyses. A) Correlation between fronto-central cluster of the P3b and the central cluster of EMSI. B) Correlation between the occipital cluster of the P3b and the central cluster of EMSI. C) Correlation between fronto-central cluster of the P3b and the centro-parietal cluster of LMSI. D) Correlation between the occipital cluster of the P3b and the centro-parietal cluster of LMSI. E) Correlation between the behavioural measure of multisensory integration and the central cluster of EMSI.

### Relating Behavioural to EEG Multisensory Integration Measures

As for the behavioural measure of multisensory integration, the only significant correlation observed was with the early EEG measure of multisensory integration (*r*(56) = .322, *p* = .014) (Figure 7E). When participants showing no significant violation were excluded from the correlation, the only significant correlation with the behavioural measure was still exclusively with the early EEG measure of multisensory integration (*r*(44) = −.406, *p* = .006). Therefore, including all participants did not change the significance of the relationship of the behavioural measure of multisensory integration with the other measures. There were no other significant correlations throughout but see Table 2 for all comparisons.

## Discussion

The purpose of this study was to determine whether a relationship exists between multisensory associative learning and multisensory integration abilities. We conducted an EEG experiment to evaluate early implicit measures of associative learning and multisensory integration, with three novel findings. First, confirming our hypothesis, we observed a significant correlation between associative learning bilaterally in fronto-central and occipital scalp areas, as indexed by the P3b, and early multisensory integration in the central scalp region. Second, this same index of multisensory associative learning was also related to the later measures of multisensory integration bilaterally in the centro-parietal scalp area. Finally, our behavioural measure of multisensory integration validated our EEG measure of multisensory integration. Our results showed that individuals who exhibited stronger neural markers of audiovisual associative learning also displayed better performance in overall integration of audiovisual information.

The most consistent observation in our data was a significant relationship between associative learning, as indexed by the P3b, and early multisensory integration. Overarchingly, this highlights the effect of higher-order processes (i.e., learned associations) in the earliest window of integration (i.e., a top-down effect). Particularly, integration was observed as early as 48 ms post stimulus presentation, and until 100 ms, which is in line with the current literature (Giard and Peronnet, 1999; Molholm, Ritter et al., 2002). Top-down effects have been previously established to have an effect, although limited, in sensory interaction prior to 100 ms (De Meo, Murray et al., 2015; Talsma, Doty et al., 2007; Talsma and Woldorff, 2005). This early index has been identified as having a centro-parietal scalp distribution (Cappe, Thut et al., 2010; Molholm, Ritter et al., 2002), which supports the current study’s findings.

The results indicate that prior learned associations may be playing a role in *how* sensory information is integrated. As the present study finds, top-down influences such as associative learning thus seem to be related to subadditive violations of the additive rule, which could reflect more efficient processing. A possible explanation for why only subadditivity was observed could be attributed to the salience of the choice of stimuli. The present study was comprised of bimodal stimuli presented at very high effectiveness, which could be responsible for activating a certain type of multisensory neuron, which have a high dynamic range and fire in an increasingly subadditive manner as stimulus effectiveness grows. (Cappe, Thut et al., 2010; Stevenson, Bushmakin et al., 2012; Perrault Jr, Vaughan et al., 2003). Furthermore, if near-ceiling effects are observed as a result of the high-salience stimuli, subadditive effects may be representative of more efficient processing as a result of the reweighting between sensory features, or rather of top-down influences such as attention (Werner and Noppeney, 2010) or, crucially, learned associations.

The P3b in both clusters is thought to be representative of inhibitory processes and of updating/encoding of the memory representation (Polich, 2007). It is worth mentioning that although we did observe a significant relationship with multisensory integration in our established time window for late associative learning, frontal activity is usually associated with P3a generation, as opposed to the typical parietal activity which is associated with the P3b. This is an important distinction, as the P3a is thought to be representative of exogenous attention-switching elicited by distractors, as opposed to memory-encoding processes by the P3b. However, there is increasing evidence highlighting the neural relationship between both components (Ebmeier, Steele et al., 1995; Soltani and Knight, 2000), which supports the notion that the relationship between bottom-up and top-down processing and their neural generators is interactive.

The later index of multisensory associative learning was also significantly correlated with the later index of multisensory enhancement exclusively in the centro-parietal cluster. As with the early measure of multisensory integration, this cluster showed subadditivity and was significantly correlated with the associative learning measures. Furthermore, the early and the late measure of multisensory integration share similar topographical profiles, which could imply that they have similar neural generators. The idea that multisensory processing possesses some level of flexibility and synchrony is becoming increasingly prevalent (Talsma, 2015) through connecting pathways between sensory cortices directly to each other (Falchier, Clavagnier et al., 2002) or through cortico-thalamic-cortical pathways (Hackett, Smiley et al., 2007; Lakatos, Chen et al., 2007; Van den Brink, Cohen et al., 2014). It is difficult to rule out that the significant relationship between associative learning in late multisensory integration is fully independent from the one in early multisensory integration. It is possible that the learned associations acted as top-down influences on the integration process as a whole. It could be stipulated, then, that later multisensory integration is independent from early integration, or rather the change in early multisensory integration could be responsible, in a downstream manner, for the multisensory integration observed later. The lack of any significant relationship between associative learning and the occipital scalp area where late multisensory enhancement was observed could be attributed to the rather low-level visual cortex activity where multisensory integration is known to occur (Foxe and Schroeder, 2005).

While the later, more attention-driven index of perceptual learning, the P3b was related to multisensory integration, the earlier, more feature-driven response, the MMN, was not related to integration. A potential reason for not seeing any effect between the early index of associative learning and overall multisensory integration could be an indication that multisensory associative learning relies on more complex higher-order processes and not simply sensory characteristics. However, it is likely that the MMN is indexing a neural process that is not related to multisensory integration

### Quantifying associative learning

In the learning phase, participants were also presented with a Deviant condition, which was different than the Mismatch condition. As expected, both the Mismatch and the Deviant conditions yielded significant MMN and P3b components. The Deviant condition was included to control for exogenous attention switching, as opposed to a detection in a deviation from the statistical pattern of shape-tone associations (Rohlf, Habets et al., 2017). As such, the infrequently-presented Mismatch pairings tended to elicit a P3b wave of lower amplitude than the Deviant stimuli, because in the latter stimuli, attention is reoriented towards the presentation of novel features themselves as opposed to the violation in pairing expectation in the Mismatch condition. The use of a three-stimulus oddball detection task was vital to providing this evidence, at the very least providing a more conservative and valid measure of differences in amplitude between the Mismatch and Match conditions. This more conservative measure is based on the fact that the Deviant stimulus is only elicited by exogenous attention switching and the lower-amplitude P3b is elicited by the Mismatch. Without the inclusion of a Deviant condition, the effect could have been difficult to isolate in the EEG signal.

### Behavioural MSI and early EEG MSI

Early neural signatures of multisensory integration in the EEG signal were significantly related to behavioural benefits in RT during a detection task. While this provides evidence that this early neural index of multisensory integration successfully captures a component of the behavioural benefits of multisensory integration, this behavioural measure did not relate to associative learning. Indeed, the magnitude of behavioural enhancement was quite small as the stimuli were very salient and were presented with no noise. The principle of inverse effectiveness explains that degraded signal from multisensory inputs result in a greater degree of multisensory gain than when the unisensory components are presented individually (Meredith and Stein, 1986). Therefore, the small multisensory behavioural benefit identified in this study is most likely as a result of including stimuli with a high signal-to-noise ratio (SNR). In this experimental design, the same novel arbitrarily-paired stimuli were used throughout this study with the purpose of preserving the validity of the measures from one phase to the other. This would ensure that any relationship between associative learning and multisensory integration that was found would be due to our experimental manipulations, and not the SNR of the stimuli themselves. We would predict that the use of less salient stimuli would result in stronger multisensory behavioural benefits, and perhaps a stronger relationship with associative learning.

### Developmental implications

These results confer many interesting developmental implications. Throughout development, there is a gradual shift towards using and relying on learned associations as opposed to solely the sensory features (e.g., timing and spatial congruence). Particular attention would be warranted when testing children in a study such as this, as they do not tend to rely on learned associations when integrating sensory information. Similarly, poor abilities in learning associations, especially from multiple sensory modalities could lead to an overreliance on stimulus features.

This phenomenon could, for example, be an issue in autistic populations, where there tends to be a bias towards processing local features over global stimulus features (Fiebelkorn, Foxe et al., 2013; Happe, 1999; Happe and Frith, 2006). It is therefore possible that populations with multisensory integration difficulties also have deficits in multisensory associative learning. For example, research in autism reveals that individuals on the spectrum show atypical looking patterns to faces (Dalton, Nacewicz et al., 2005; Spezio, Adolphs et al., 2007; Stevenson, Philipp-Muller et al., 2019; Trepagnier, Sebrechts et al., 2002), and also show decreased multisensory integration (Baum, Stevenson et al., 2015; Feldman, Dunham et al., 2018; Stevenson, Philipp-Muller et al., 2019) opening the possibility that a lack of exposure to the visual components of speech (e.g., the lips moving and mouthing the syllables) is related to poorer performance in multisensory integration. This could in turn play a key role in the reason why individuals in this population tend to have an overreliance on the sensory cues to bind (i.e., spatial and temporal congruence), as opposed to a balanced re-weighting between stimulus features and learned associations.

### Limitations and future directions

Future studies should parametrically manipulate the choice of stimuli to include stimuli that have a lower SNR. This would be key in determining the extent of the relationship between learned associations and multisensory integration, insofar as stimulus manipulations allow. Furthermore, studies including more ecologically-valid higher-level stimuli, such as multisensory speech, could be useful in extending the generalizability of the important relationship between associative learning and multisensory integration. Furthermore, future studies could attempt to maximize multisensory associative learning at different developmental stages, which already is showing some promising results (Rohlf, Habets et al., 2017). These studies could also test for multisensory integration with the use of the learned associations to see what could be modulating performance for multisensory integration. Furthermore, these studies could investigate further into how these relationships changed across age groups.

The present study was able to establish a direct link between associative learning and the capacity to integrate information from multiple sensory modalities. Participants who showed stronger indices of associative learning also exhibited stronger indices of multisensory integration of the stimuli they learned to associate. Specifically, fronto-central and occipital scalp areas exhibiting significant P3b signatures were significantly correlated with central scalp areas showing neural signatures of early integration and one centro-parietal scalp area showing later multisensory integration. Furthermore, our behavioural index was significantly related to our early measure of multisensory integration, thus serving as a validation for our measure. This study highlights the key influence of top-down effects such as multisensory associative learning on multisensory integration.

## Acknowledgements

RS is funded by an NSERC Discovery Grant (RGPIN-2017-04656), a SSHRC Insight Grant (R5502A07), the University of Western Ontario Faculty Development Research Fund, the province of Ontario Early Researcher Award, and a Canadian Foundation for Innovation John R. Evans Leaders Fund (37497).

## Notes

### Competing Interest Statement

The authors have declared no competing interest.

